# Ciliary ARL13B inhibits developmental kidney cystogenesis in mouse

**DOI:** 10.1101/2023.02.08.527739

**Authors:** Robert E. Van Sciver, Alyssa B. Long, Harrison G. Katz, Eduardo D. Gigante, Tamara Caspary

## Abstract

ARL13B is a small GTPase enriched in cilia. Deletion of *Arl13b* in mouse kidney results in renal cysts and an associated absence of primary cilia. Similarly, ablation of cilia leads to kidney cysts. To investigate whether ARL13B functions from within cilia to direct kidney development, we examined kidneys of mice expressing an engineered cilia-excluded ARL13B variant, ARL13B^V358A^. These mice retained renal cilia and developed cystic kidneys. Because ARL13B functions as a guanine nucleotide exchange factor (GEF) for ARL3, we examined kidneys of mice expressing an ARL13B variant that lacks ARL3 GEF activity, ARL13B^R79Q^. We found normal kidney development with no evidence of cysts in these mice. Taken together, our results show that ARL13B functions within cilia to inhibit renal cystogenesis during mouse development, and that this function does not depend on its role as a GEF for ARL3.

## Introduction

Primary cilia are found on nearly every cell in vertebrates and are linked to cell signaling. These solitary microtubule-based appendages are required for normal kidney development (Calvet, 2002; Pazour, 2004). In mice and zebrafish, constitutive ablation of cilia leads to kidney cysts (Pazour et al., 2000; Sun et al., 2004; Yoder et al., 2002). Patients with mutations in ciliary proteins exhibit renal anomalies including cysts (Doherty, 2009; Hildebrandt et al., 2011; Lehman et al., 2008). Critical signaling receptors such as the polycystin proteins PC1 and PC2, whose loss leads to renal cystogenesis, localize to renal primary cilia (Pazour et al., 2002; Yoder et al., 2002). Mice carrying a mutation in PC2 that precludes its cilia localization develop renal cysts indistinguishable from mice lacking renal PC2 (Walker et al., 2019; Wu et al., 1998). Such evidence points to ciliary signaling playing an important role in kidney development; however, the molecular details specific for renal ciliary signaling during development remain murky.

ARL13B is an ADP Ribosylation Factor Like (ARL) GTPase highly enriched in cilia. ARL13B was discovered in a zebrafish forward genetic screen for renal cystic phenotypes (Sun et al., 2004). Null mutations in mouse *Arl13b* are embryonic lethal prior to kidney development (Caspary et al., 2007; Su et al., 2012). ARL13B’s GTPase domain contains the canonical switch 1 and switch 2 and N-terminal helix required for interaction with putative effectors (Kahn et al., 2014; Mariani et al., 2016; Miertzschke et al., 2013). ARL13B also possesses some unusual features for an ARL protein. ARL13B is predicted to use an atypical GTPase cycle, since it has a glycine (G75) where almost all other GTPases have glutamine, an amino acid residue that is directly involved in GTP hydrolysis (Ivanova et al., 2017). In addition to the GTPase domain, it contains a 20 kDa novel C-terminal domain which includes a VxPx motif required for ARL13B cilia localization (Mariani et al., 2016). ARL13B functions as a guanine nucleotide exchange factor (GEF) for another ARL protein, ARL3, which also localizes to cilia (Zhou et al., 2006). As a GEF, ARL13B facilitates the exchange of GTP for GDP, activating ARL3. In the absence of ARL13B, ARL3 cannot be detected in cilia, possibly because ARL3’s activation is needed for its ciliary retention (Gigante et al., 2020; Gotthardt et al., 2015). Additionally, ARL13B regulates the phospholipid composition of the ciliary membrane via inositol polyphosphate-5-phosphatase E (INPP5E) (Humbert et al., 2012). Normally, INPP5E catalyzes the hydrolysis of the 5-phosphate from PI(4,5)P_2_, maintaining PI(4)P levels on the ciliary membrane (Bielas et al., 2009; Garcia-Gonzalo et al., 2015; Jacoby et al., 2009). Phospholipids on the ciliary membrane regulate ciliary signaling by controlling cilia-localized proteins (Nachury et al., 2007; Santagata et al., 2001). For example, TULP3 is a PI(4,5)P_2_-binding protein important for trafficking ciliary components into the cilium (Mukhopadhyay et al., 2010; Norman et al., 2009). In mice, mutations in *Arl13b, Arl3, Inpp5e* or *Tulp3* during development cause renal cysts (Hakim et al., 2016; Hwang et al., 2019; Legue and Liem, 2019; Li et al., 2016; Schrick et al., 2006; Seixas et al., 2016).

Conditional deletion of *Arl13b* in mouse kidney leads to renal cysts in all regions of the nephron (Li et al., 2016; Seixas et al., 2016). These *Arl13b*-null kidneys lack cilia altogether. Since the loss of renal cilia can also lead to cysts (Davenport et al., 2007; Jonassen et al., 2008; Lin et al., 2003; Yoder et al., 1996), it is not possible to decipher whether kidney-specific deletion of *Arl13b* causes cysts because of loss of ARL13B function or loss of cilia (Li et al., 2016; Seixas et al., 2016). The loss of cilia in ARL13B-deleted kidneys is unusual. When ARL13B is deleted in the mouse embryonic node, neural tube or fibroblasts derived from *Arl13b*-null embryos, cilia are present albeit short (Caspary et al., 2007; Dilan et al., 2019; Hanke-Gogokhia et al., 2017; Larkins et al., 2011; Lu et al., 2015). The necessity of ARL13B for renal cilia suggests the possibility that ARL13B functions differently in kidney tissue compared to other cell types. Potentially relevant to this, the PI(4,5)P_2_-binding protein TULP3 is essential for ciliary localization of ARL13B in kidney but not in other cell types (Ferent et al., 2019; Hwang et al., 2019; Legue and Liem, 2019; Palicharla et al., 2023). Thus, it is important to dissect discrete functions of ARL13B in kidney.

Here, we address two questions concerning ARL13B function in kidney development using mouse alleles carrying specific point mutations at the endogenous locus. First, we ask whether ARL13B functions from within cilia to regulate renal development. We examine kidney development in mice expressing an ARL13B variant that that does not localize to cilia due to a V358A mutation in the VxPx motif required for ARL13B ciliary localization (Gigante et al., 2020; Mariani et al., 2016). Second, we ask whether ARL13B’s function in renal development requires its GEF activity for ARL3 in renal development. We analyze kidneys of mice expressing a variant that lacks ARL13B GEF activity for ARL3 due to an R79Q mutation (Ivanova et al., 2017; Suciu et al., 2021). Engineering the mutations at the endogenous *Arl13b* locus enables us to unravel ARL13B’s function in renal development under physiological conditions.

## Results

### *Arl13b*^*V358A/V358A*^ mice develop heavier kidneys that lack ciliary ARL13B

Although complete deletion of ARL13B in mice is lethal prior to kidney development, conditional deletion of ARL13B in the kidneys leads to cysts and a loss of cilia (Li et al., 2016; Seixas et al., 2016; Su et al., 2012). In contrast, mice expressing ARL13B^V358A^ or ARL13B^R79Q^ survive into adulthood (Gigante et al., 2020; Suciu et al., 2021). To assess the effect of these mutations in the kidney, we weighed the kidneys of 18-week-old animals. We observed a significant increase in total kidney weight of male and female homozygous *Arl13b*^*V358A/V358A*^ mice compared to sex-matched heterozygous *Arl13b*^*V358A/+*^ and wild type control animals (Fig 1A). In contrast, the kidneys of *Arl13b*^*R79Q/R79Q*^ mice weighed the same as wild type control animals. We also analyzed these data as kidney weight:body weight ratios and found the same result (Fig S1). These data suggest loss of ciliary ARL13B impacts kidney weight.

**Figure 1.**
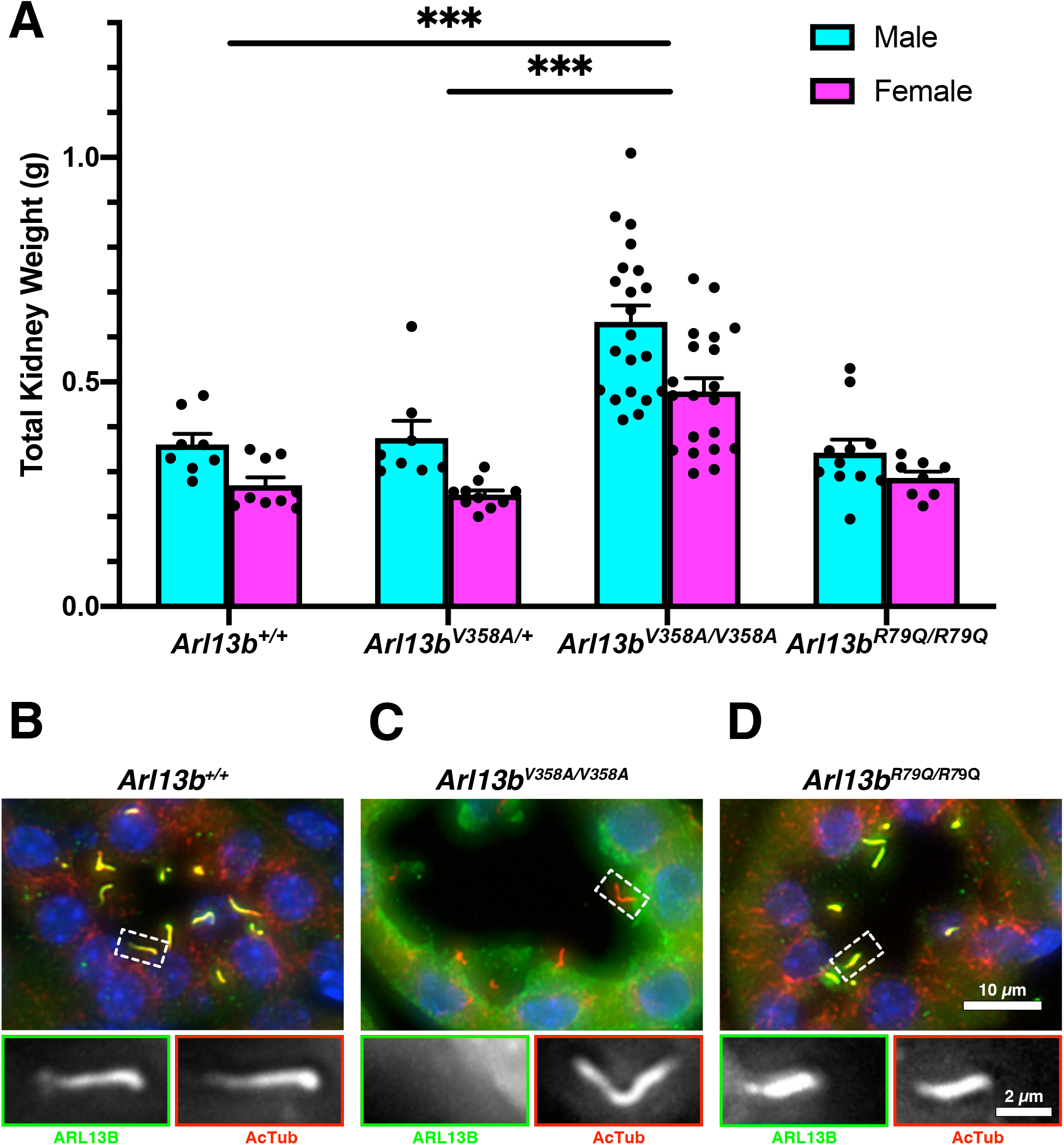
*Arl13b*^*V358A/V358A*^ mice develop heavier kidneys that lack ciliary ARL13B. (A) Quantification of the total kidney weight for kidneys from 18-week-old male and female mice, mean ± SEM. n ≥ 8 per genotype and sex. Two-way ANOVA, *** p < 0.001. (B-D) Kidney sections stained with antibodies against ARL13B (green) and acetylated α-tubulin (red). Scale bar for large panels: 10 μm; scale bar for insets: 2 μm.

To evaluate the subcellular localization of ARL13B in the kidneys of these mutant mice, we performed immunofluorescent staining for ARL13B and acetylated α-tubulin, a marker for primary cilia. In wild type kidneys, ARL13B and acetylated α-tubulin colocalized in primary cilia (Fig 1B). In *Arl13b*^*V358A/V358A*^ kidneys, we found acetylated α-tubulin positive cilia that lacked ARL13B staining, consistent with previous findings in other tissue types (Fig 1C) (Gigante et al., 2020; Guo et al., 2017; Higginbotham et al., 2012; Mariani et al., 2016). In *Arl13b*^*R79Q/R79Q*^ kidneys, we observed ARL13B colocalized with acetylated α-tubulin in line with previous work (Fig 1D) (Humbert et al., 2012; Mariani et al., 2016). Taken together, these data indicate that ARL13B^V358A^ and ARL13B^R79Q^ endogenous expression in kidney aligns with what is observed in other cell types (Gigante et al., 2020; Guo et al., 2017; Higginbotham et al., 2012; Humbert et al., 2012; Mariani et al., 2016; Roy et al., 2017).

### *Arl13b*^*V358A/V358A*^ mice develop cystic kidneys, whereas *Arl13b*^*R79Q/R79Q*^ mice do not

To examine the enlarged kidneys more closely, we performed hematoxylin and eosin (H&E) staining of paraffin sections. We examined *Arl13b*^*R79Q/R79Q*^ kidneys at multiple stages of development. The *Arl13b*^*R79Q/R79Q*^ kidneys appeared morphologically normal through 18 weeks of age and we did not detect any dilations or cysts (Fig 2A). In contrast, we observed tubule dilations in *Arl13b*^*V358A/V358A*^ kidneys starting at postnatal day (P) 10 and cysts at weaning age (P24), with large cysts detected at ∼4 months (P120; Fig 2B). We analyzed H&E-stained kidneys from 18-week-old animals and observed a cystic index (% cystic area divided by total kidney area) of 9.48±1.51% and 7.57±1.03% in male and female control mice, respectively. We observed a cystic index of 29.97±5.28% and 30.41±6.24% in male and female *Arl13b*^*V358A/V358A*^ mice, respectively (Fig 2C). Thus, we found a higher cystic index when ARL13B was excluded from cilia in both male and female kidneys compared to control kidneys

**Figure 2.**
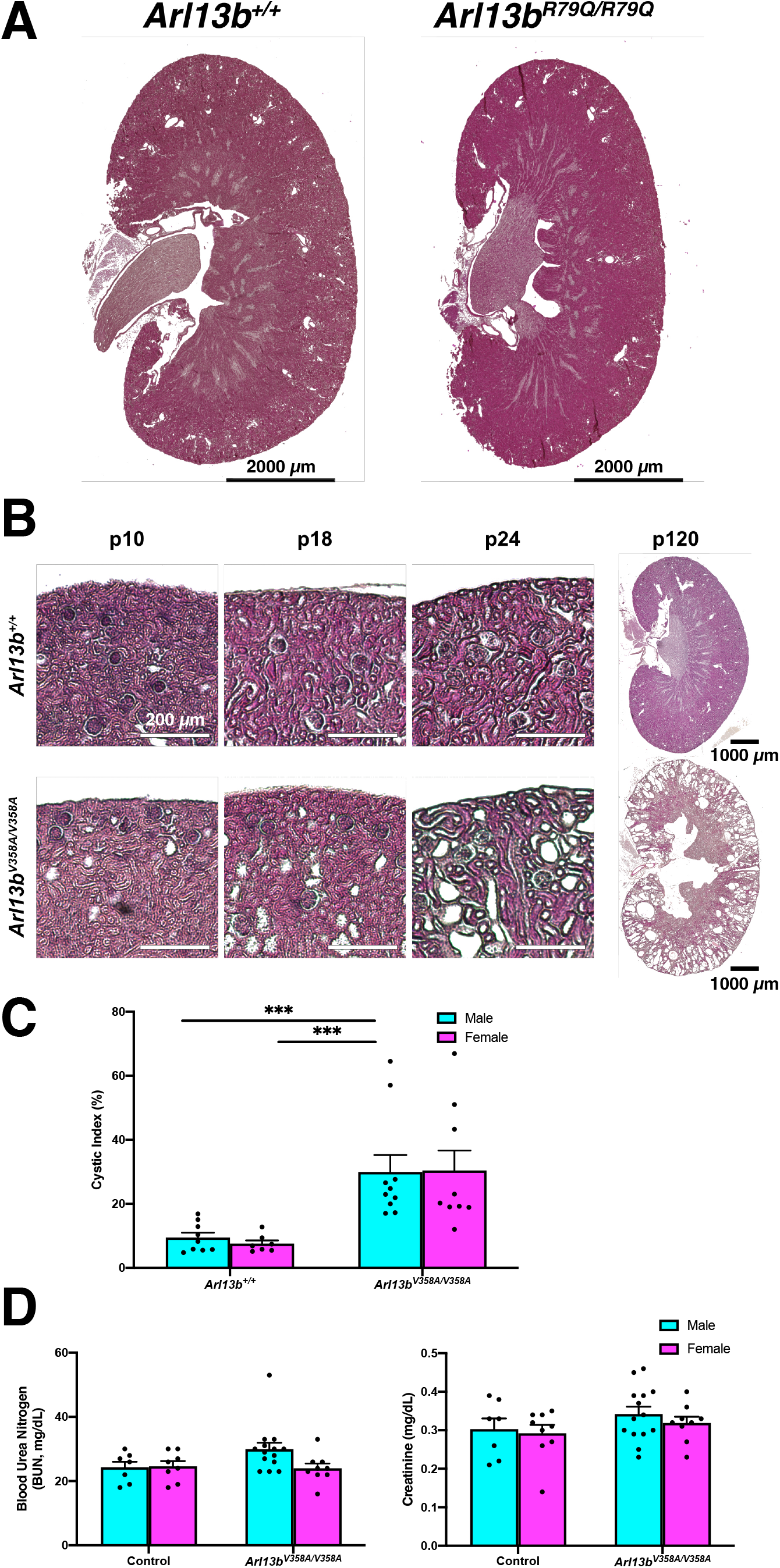
*Arl13b*^*V358A/V358A*^ mutant mice exhibit a progressive cystic kidney phenotype. (A) H&E staining of kidney cross sections from 18-week-old male mice. Scale bar: 2000 μm. (B) Tubule dilations appeared in *Arl13b*^*V358A/V358A*^ kidneys around P10 and progressed to cysts by weaning age (P24). Scale bars in P10, P18, P24: 200 μm. Scale bar in P120: 1000 μm. (C) Cyst formation in *Arl13b*^*V358A/V358A*^ quantified by cystic index, mean ± SEM. n ≥ 7 mice per genotype and sex. Two-way ANOVA, ** p < 0.01. (D) *Arl13b*^*V358A/V358A*^ mice display normal renal physiology: Blood urea nitrogen (BUN) and serum creatinine levels were measured from 18-week old male and female mice, mean ± SEM. n ≥ 7 per genotype and sex. Two-way ANOVA, no significant change between *Arl13b*^*V358A/V358A*^ and control animals.

To assess kidney function, we performed physiological analysis on serum from 18-week-old animals and measured blood urea nitrogen (BUN) and creatinine levels. We did not detect significant changes in these measures in *Arl13b*^*V358A/V358A*^ mice of either sex compared with control mice (Fig 2D). Taken together these data indicate that *Arl13b*^*V358A/V358A*^ mice develop progressive cystic kidneys with normal kidney physiology. Furthermore, these data indicate that the kidney phenotype in the *Arl13b*^*V358A/V358A*^ mice does not depend on the GEF activity of ARL13B.

### Cysts in *Arl13b*^*V358A/V358A*^ kidneys are detected in all regions of the nephron

To investigate the *Arl13b*^*V358A/V358A*^ cysts, we analyzed where they form within the nephron by looking at multiple segment-specific markers in kidney sections. We stained sections with *Lotus tetragonobolus* lectin (LTL) and *Dolichos biflorus* agglutinin (DBA) to identify proximal tubules and collecting ducts, respectively, and found cysts in both nephron segments (Fig 3A and B). We also stained sections with antibody to sodium chloride cotransporter (NCC), labeling the distal convoluted tubule, and observed cysts in this nephron region (Fig 3C, cyan). Lastly, we stained sections with antibody to Tamm-Horsfall protein (THP), or uromodulin, which revealed cysts in the thick ascending limb of the loop of Henle (Fig 3C, magenta). We noted uromodulin-positive crystal deposits, which are implicated in cyst pathogenesis, in this region. These findings indicate that the *Arl13b*^*V358A/V358A*^ mutation leads to cysts in all segments of the nephron.

**Figure 3.**
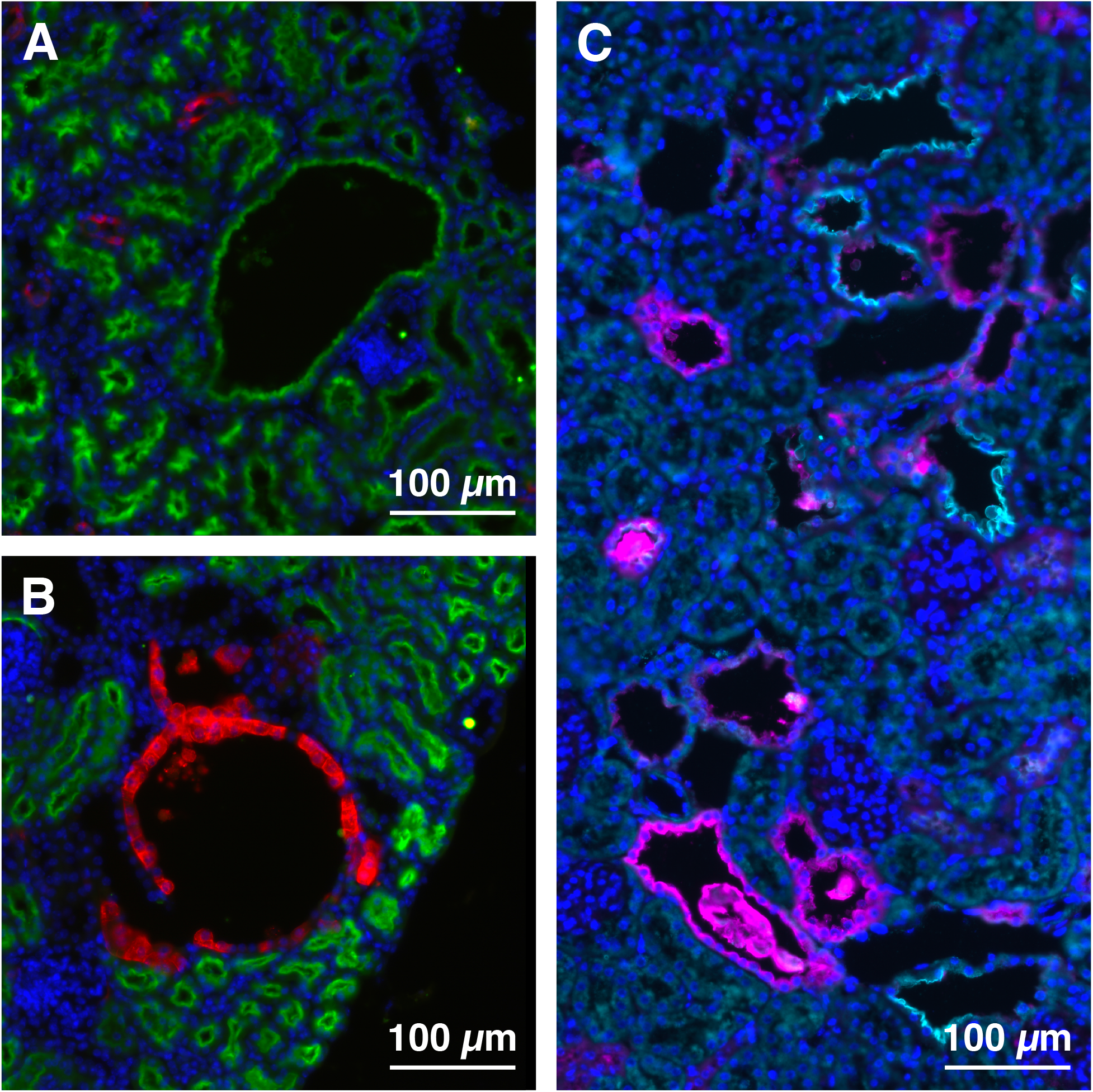
Cysts in *Arl13b*^*V358A/V358A*^ kidneys are present in all nephron segments. (A) LTL (proximal tubule, green), (B) DBA (collecting duct, red), (C) THP (thick ascending limb of the loop of Henle, magenta) and NCC (distal convoluted tubule, cyan) staining in *Arl13b*^*V358A/V358A*^ kidney sections. Scale bars: 100 μm.

### Cilia length is normal in *Arl13b*^*V358A/V358A*^ kidneys, but ciliation rate is reduced in cystic regions

Our previous work showed that immortalized fibroblasts derived from *Arl13b*^*V358A/V358A*^ embryos have fewer and shorter cilia than wild type (Gigante et al., 2020). To test whether cilia were altered in *Arl13b*^*V358A/V358A*^ kidneys, we measured cilia length using acetylated α-tubulin. In wild type animals, cilia of the renal epithelium were 2.436±0.136 μm. In *Arl13b*^*V358A/V358A*^ animals, cilia of normal tubule renal epithelium were 2.522±0.194 μm whereas cilia of the renal epithelia lining cystic regions were 2.755±0.213 μm (Fig 4A). These measurements indicate that the loss of ciliary ARL13B does not affect cilia length of the renal epithelial cells of the normal tubules or cystic regions in *Arl13b*^*V358A/V358A*^ kidneys. We next asked whether there was a change in ciliation rate as measured by the number of cilia per centriolar pair. We stained kidney sections with acetylated α-tubulin and fibroblast growth factor receptor 1 oncogene partner (FGFR1OP, FOP) to stain cilia and centriolar satellites, respectively. In wild type kidney sections, we observed a ciliation rate of 94.00±5.13% (Fig 4B). In *Arl13b*^*V358A/V358A*^ kidneys, we found an average ciliation rate of 81.33±5.812% in normal tubules and 60.67±6.69% in cystic epithelia (Fig 4B). The ciliation rate in *Arl13b*^*V358A/V358A*^ cystic tubules was not significantly different from that in *Arl13b*^*V358A/V358A*^ normal tubules but was significantly different from the ciliation rate in wild type tubules. Taken together, these data reveal that the majority of cells in *Arl13b*^*V358A/V358A*^ kidneys are ciliated and display cilia of normal length.

**Figure 4.**
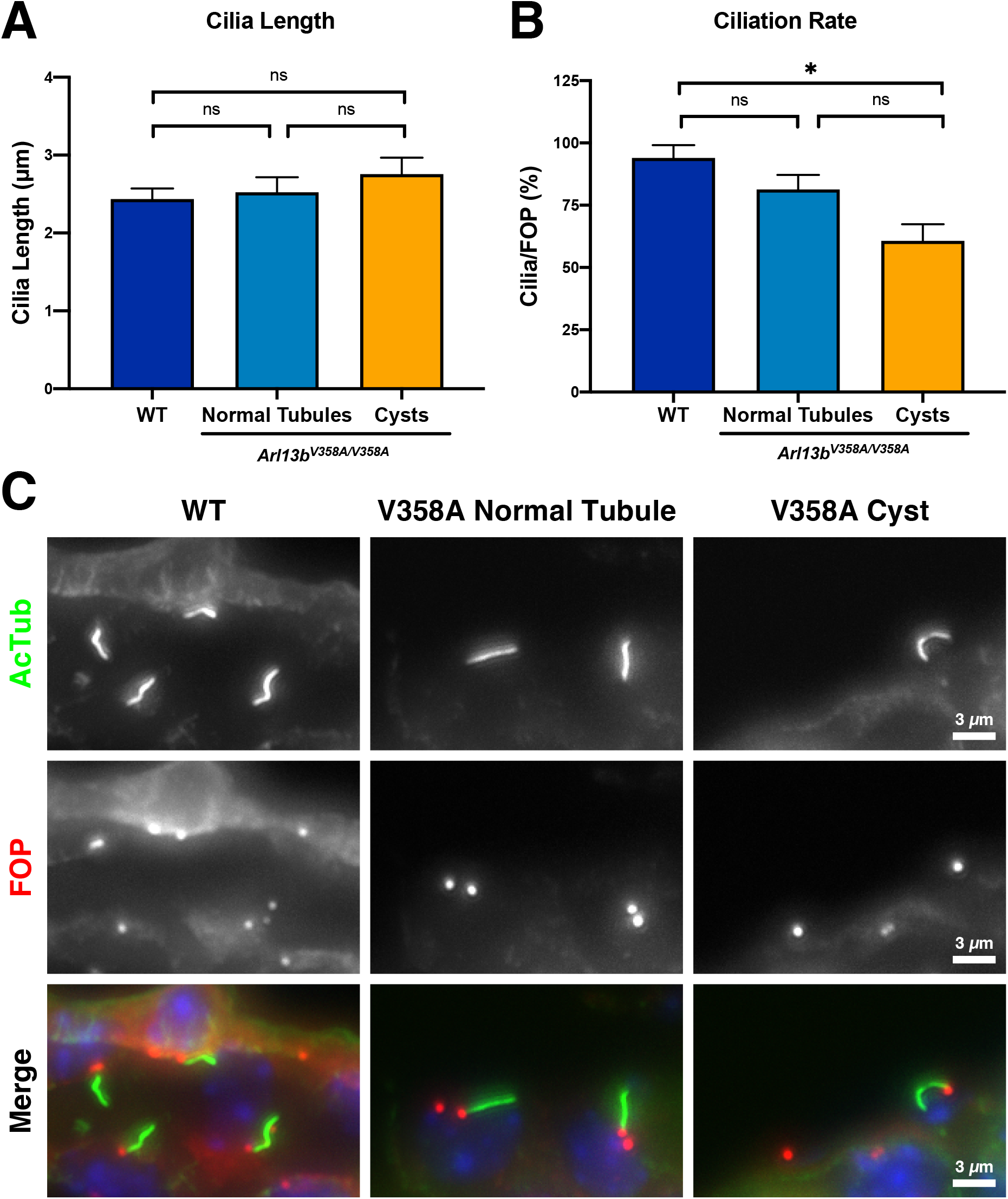
Cilia length is normal in *Arl13b*^*V358A/V358A*^ kidneys, but ciliation rate is reduced in *Arl13b*^*V358A/V358A*^ cystic regions. (A) Quantitation of cilia length in wild type kidneys as well as normal tubules and cystic regions of *Arl13b*^*V358A/V358A*^ kidneys, mean ± SEM. (B) Quantification of ciliation rate (cilia per basal body as indicated by acetylated alpha-tubulin and FGFR1OP (FOP) staining) in wild type kidneys and normal tubules and cystic regions of *Arl13b*^*V358A/V358A*^ kidneys, mean ± SEM. n = 3 mice per genotype with 100+ cilia counted per animal. One-way ANOVA, * p < 0.05. (C) Representative staining of acetylated tubulin and FOP in wild type kidneys and normal and cystic regions of *Arl13b*^*V358A/V358A*^ kidneys. Scale bars: 3 μm.

### Ciliary exclusion of ARL13B affects the localization patterns of ARL3 and TULP3

To explore the mechanism of ARL13B action within cilia, we investigated the ciliary localization of ARL13B effectors, interactors and proteins involved in cystogenesis: ARL3, TULP3, PC2 and cystin (Cai et al., 1999; Hwang et al., 2019; Legue and Liem, 2019; Schrick et al., 2006; Tao et al., 2009). We found that ARL3 localized to cilia of wild type and *Arl13b*^*R79Q/R79Q*^ mutant kidneys but was virtually undetectable in cilia of *Arl13b*^*V358A/V358A*^ kidneys (0.086±0.011 compared to control, Figure 5A’-A’’’’). TULP3 is a PI(4,5)P_2_-binding protein so its ciliary localization reflects ciliary PI(4,5)P_2_ levels (Garcia-Gonzalo et al., 2015). We were unable to detect TULP3 in wild type and *Arl13b*^*R79Q/R79Q*^ kidney cilia; however, we observed a six-fold increase in ciliary TULP3 in *Arl13b*^*V358A/V358A*^ mutant kidney cilia (5.99±0.51 compared to control, Fig 5B’-B’’’’). Together, these data are consistent with ARL13B’s known regulation of INPP5E ciliary localization: without ciliary ARL13B, ciliary INPP5E levels decrease and PI(4,5)P_2_ levels increase, causing ciliary enrichment of TULP3 (Garcia-Gonzalo et al., 2015).

**Figure 5.**
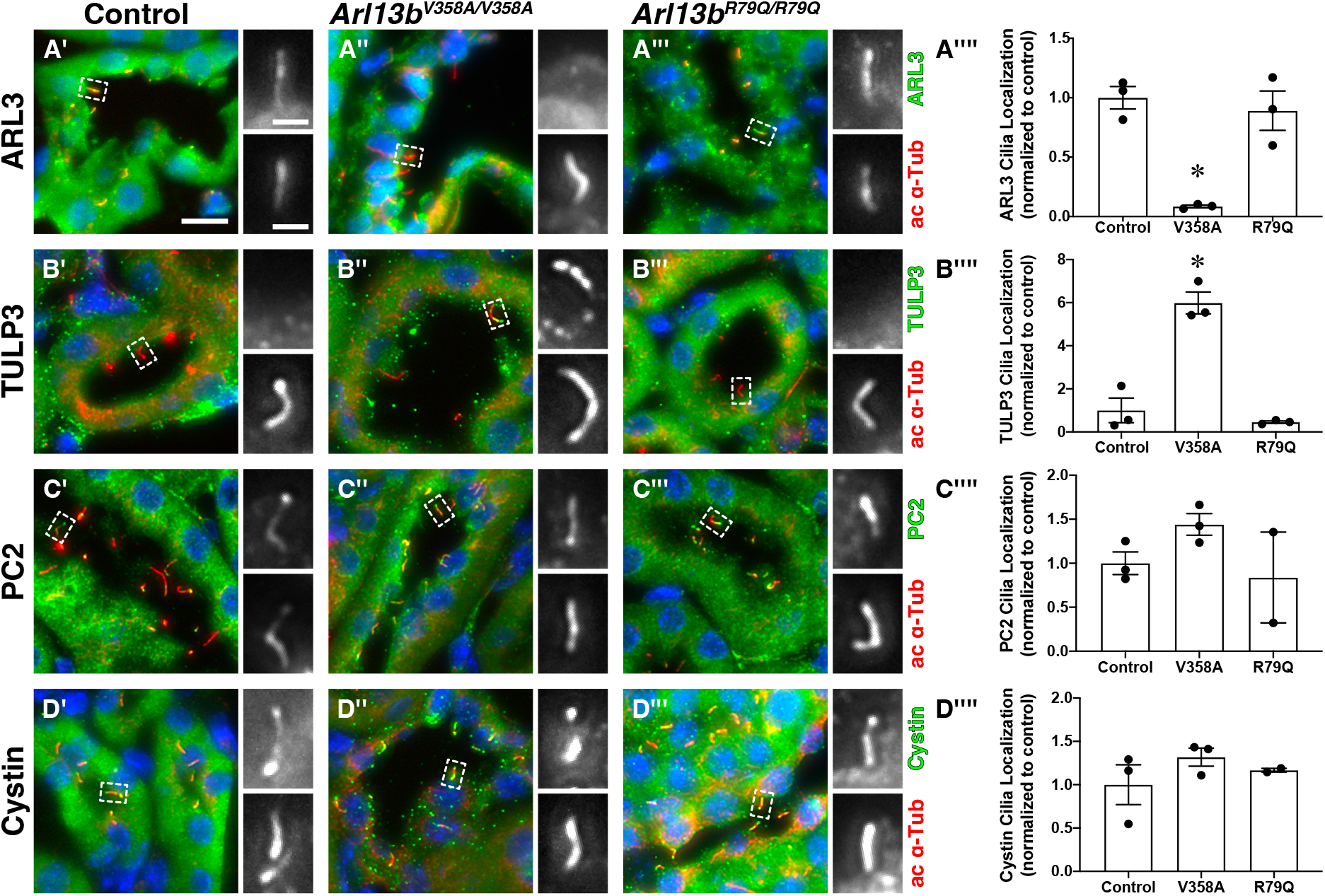
Ciliary exclusion of ARL13B affects the localization patterns of ARL3 and TULP3. Acetylated α-tubulin (ac α-Tub, red, lower panel of insets) stains cilia. (A’, A’’, A’’’) ARL3 (green, upper panel of insets) is in cilia of control and *Arl13b*^*R79Q/R79Q*^ kidneys. (B’, B’’, B’’’) TULP3 (green, upper panel of insets) is localized to cilia in *Arl13b*^*V358A/V358A*^ kidneys. (C’, C’’, C’’’) Polycystin 2 (PC2, green, upper panel of insets) and (D’, D’’, D’’’) cystin (green, upper panel of insets) are normally localized in both *Arl13b*^*V358A/V358A*^ and *Arl13b*^*R79Q/R79Q*^ kidneys. Scale bar for large panels: 10 μm; scale bar for insets: 2 μm. (A’’’’, B’’’’, C’’’’, D’’’’) Quantification of cilia localization (cilia positive for a given stain, normalized to control animals) in control, *Arl13b*^*V358A/V358A*^, and *Arl13b*^*R79Q/R79Q*^ kidneys, mean ± SEM. An average of 92 cilia were counted per stain per animal. n = 3 mice per genotype per stain, except for PC2- and cystin-stained *Arl13b*^*R79Q/R79Q*^ kidneys, n = 2. One-way ANOVA, * p < 0.05 vs control.

We next looked at two proteins with known roles in cystogenesis, polycystin 2 (PC2) and cystin. We observed ciliary PC2 and cystin in wild type, *Arl13b*^*V358A/V358A*^, and *Arl13b*^*R79Q/R79Q*^ kidney sections (Fig 5C’-C’’’’, D’-D’’’’). These results indicate that PC2 and cystin localization do not depend on ciliary ARL13B or its GEF activity.

## Discussion

Our findings indicate that ARL13B activity, specifically in cilia, normally suppresses cystogenesis during development, and that ARL13B GEF activity for ARL3 is dispensable for this process. Using point mutations targeted to the endogenous *Arl13b* locus, we analyzed the consequence of disrupting cilia localization of ARL13B (ARL13B^V358A^) or of loss of ARL13B GEF activity for ARL3 (ARL13B^R79Q^). While we did not observe renal cysts in *Arl13b*^*R79Q/R79Q*^ kidneys, we found normal cilia in *Arl13b*^*V358A/V358A*^ kidneys along with progressive cystogenesis. Taken together, these data suggest that ARL13B functions from within cilia to inhibit cysts and implies ARL13B’s GTPase function, but not its GEF activity, is important for this role.

These experiments resolve a previously unanswered question in the field. Conditional deletion of *Arl13b* leads to renal cysts but also a complete loss of cilia (Li et al., 2016; Seixas et al., 2016). Thus, despite the enrichment of ARL13B in cilia, it was unclear whether ARL13B functioned there, as loss of cilia was known to result in renal cysts. ARL13B^V358A^ protein is undetectable in kidney cilia yet retains the known biochemical activities of ARL13B, so the renal cysts in *Arl13b*^*V358A/V358A*^ kidneys argue that ARL13B functions from within cilia (Gigante et al., 2020; Mariani et al., 2016). That said, the kidney cystogenesis arising from complete *Arl13b* loss is more severe than what we observed in *Arl13b*^*V358A/V358A*^ kidneys, raising the possibility that ARL13B may play additional functional roles in regulating kidney cysts from non-ciliary (cellular) locales (Li et al., 2016; Seixas et al., 2016). As found for every other ARL protein examined, ARL13B likely functions in multiple locales in cells in concert with different regulators and effectors; these interactions can be transient making some subcellular localizations challenging to detect (Jian et al., 2010; Nawrotek et al., 2016; Sztul et al., 2019). It is currently not clear where within the renal cells ARL13B localizes. Previous reports have detected it in early endosomes and dorsal circular ruffles in other cell types (Barral et al., 2012; Casalou et al., 2014).

The presence of normal cilia in *Arl13b*^*V358A/V358A*^ kidneys highlights the distinct role ARL13B appears to play in kidney. In other tissues, loss of ARL13B leads to short cilia, even when only excluded from cilia (Caspary et al., 2007; Dilan et al., 2019; Gigante et al., 2020; Hanke-Gogokhia et al., 2017; Larkins et al., 2011; Lu et al., 2015). Thus, renal cilia do not require ciliary ARL13B function for ciliogenesis as other cell types do, consistent with growing evidence for distinct cilia in different cell types (Kiesel et al., 2020; Silva et al., 2017; Sun et al., 2019; Zabeo et al., 2019). It remains unclear whether ARL13B plays any role in renal cilia maintenance and cyst appearance may precede the loss of renal cilia in *Arl13b*^*V358A/V358A*^ kidneys. Non-cystic regions of *Arl13b*^*V358A/V358A*^ kidneys displayed a normal number of cilia per centriolar pair, and we only observed a reduction in ciliation of cystic regions. Further experiments are needed to establish the temporal order of cyst formation and cilia loss.

ARL13B localization in renal cilia requires TULP3, which is not the case in other cell types (Ferent et al., 2019; Hwang et al., 2019; Legue and Liem, 2019; Palicharla et al., 2023). Previous work on TULP3 proposed that it regulates a cilia-specific pathway that normally inhibits cystogenesis, termed the cilia localized cyst inhibitory (CLCI) pathway (Walker et al., 2022). Our findings here indicate that ciliary ARL13B is a likely component of the CLCI pathway during mouse development. The molecular mechanism of this pathway remains unclear. While *Tulp3* mutants display a reduction of ciliary polycystin 2, we found normal ciliary enrichment of polycystin 2 and cystin proteins in *Arl13b*^*V358A/V358A*^ renal cilia (Hwang et al., 2019; Legue and Liem, 2019). Furthermore, despite previous work linking ARL13B and vertebrate Hedgehog signaling, ARL13B^V358A^ mediates Hh signaling normally (Caspary et al., 2007; Gigante et al., 2020; Larkins et al., 2011; Mariani et al., 2016). The increased localization of TULP3 in *Arl13b*^*V358A/V358A*^ renal cilia suggests the pathway is sensitive to ciliary PI(4,5)P_2_ levels. As these are regulated by INPP5E, we predict INPP5E would not be retained in *Arl13b*^*V358A/V358A*^ renal cilia. This would be consistent with data from *Arl13b*^*V358A/V358A*^ MEFs and *ARL13B* knockout RPE1 cells (Gigante et al., 2020; Qiu et al., 2021), however we await the development of antibodies that work well on tissue sections to test this hypothesis.

INPP5E also requires activated ARL3 for its ciliary targeting in other cell types, therefore we predicted a role for ARL13B’s ARL3 GEF activity in cystogenesis. Thus, we were a bit surprised that GEF-deficient *Arl13b*^*R79Q/R79Q*^ mice did not exhibit renal cystic phenotypes and ARL3 remained detectable in the renal cilia of *Arl13b*^*R79Q/R79Q*^ mice. At one level, this is consistent with the Joubert patient carrying the *ARL13B*^*R79Q*^ mutation not presenting with renal phenotypes (Cantagrel et al., 2008). However, *ARL13B*^*R79Q*^ does not rescue the cystic phenotype observed in *Arl13b*^*sco*^ mutant zebrafish (Cantagrel et al., 2008). It is unclear whether this discrepancy is due to species differences or experimental differences such as level or timing of expression. It is also not clear whether the ARL3 detected in cilia is activated. If ciliary ARL13B inhibits renal cystogenesis via ARL3, one possibility is that ARL3 may be activated by an alternative, unknown GEF. Recent work implicates RABL2 as a GEF for ARL3 in *Chlamydomonas reinhardtii*, although whether this activity is conserved in vertebrates or tissues such as the kidney remains to be determined (Liu et al., 2022; Zhang et al., 2022). Of note, GEF activities in general and also true for ARL3 GEF activity, are increasingly being found to be subject to factors that are incompletely understood or characterized. These include phospholipids, “co-GEFs” and effectors; in a small space like the cilium, they may also include concentrations of nucleotides, calcium or pH. Within cilia, such factors may be regulated by ciliary ARL13B. For example, ciliary ARL13B normally retains INPP5E on the ciliary membrane, so the loss of ciliary ARL13B is expected to indirectly alter ciliary phospholipid (PIPs) composition, with other potential indirect consequences to ciliary biology. This is consistent with ciliary enrichment of PI(4,5)P_2_-sensitive TULP3 in *Arl13b*^*V358A/V358A*^ kidneys; however, we cannot rule out residual INPP5E function.

In summary, we provide in vivo evidence that ciliary ARL13B functions to inhibit renal cystogenesis during mouse development. The relationship of cilia and ciliary signaling to renal cysts changes precipitously over time. Indeed, genetic ablation of cilia during development leads to cystogenesis whereas such deletion after postnatal day 14 leads to mild, slow-growing cysts (Davenport et al., 2007; Lin et al., 2003; Piontek et al., 2007). Whether the ciliary role of ARL13B in inhibiting cystogenesis changes over time is unknown. As a GTPase, ARL13B undoubtedly works through specific effectors in different places at distinct times. Our data show that ARL13B functions during development to inhibit kidney cystogenesis from within renal cilia and independent of its GEF activity.

## Materials and methods

### Mouse lines

All mice were cared for in accordance with NIH guidelines and Emory University’s Institutional Animal Care and Use Committee (IACUC). Lines used were *Arl13b*^*V358A*^ (C57BL/6J-*Arl13b*^*em1Tc*^) [MGI: 6256969], and *Arl13b*^*R79Q*^ (C57BL/6J-*Arl13b*^*em2Tc*^) [MGI: 6279301] (Gigante et al., 2020; Suciu et al., 2021). Mice were genotyped for the V358A mutation using primers MB11: 5’-CCTATATTCTTCTAGAAAACAGTAAGAAGAAAACCAAGAAACTAAGACTCCTTTTCATTCATC GGGC -3’ and MB12: 5’-GACAGTAAAGGATTCTTCCTCACAACCTGAC-3’ to detect the mutant allele, and primers MB21: 5’-CTTAAGATGACTTTGAGTTTGGAAGAAATACAAGATAGC-3’ and MB22: 5’-GCGTGGGACTCTTTGGAGTAGACTAGTCAATACAGACGGGTTCTA-3’ to detect the wildtype allele. Band sizes were 395bp for wildtype and 273bp for mutant. Mice were genotyped for the R79Q mutation using primers 223_2F: 5’-TCACTTGCAACAGAGCATCC-3’ and 223_2R: 5’-ACAGCTCTGCCCGTGTTTAC-3’ followed by Sanger sequencing the 304bp PCR product. All animals in this study were analyzed at 18-weeks, except as indicated in Figure 2.

### Tissue and blood harvesting

Mice were euthanized at the indicated ages by isoflurane inhalation and weighed before perfusion. Blood was harvested by cardiac puncture, followed by perfusion with ice-cold PBS and ice-cold 4% paraformaldehyde. Kidneys were harvested and weighed following perfusion. Kidneys were bisected sagittally, with one half prepared for paraffin embedding and the other half prepared for cryo-embedding. For paraffin embedding, tissues were dehydrated at room temperature for one hour each through 70% ethanol, 90% ethanol, three changes of 100% ethanol, followed by two 1 hour washes with SafeClear (Fisher). Dehydrated kidneys were placed into paraffin at 60°C overnight, then transferred to fresh paraffin at 60°C for 1 hour before embedding. For cryo-embedding, tissues were incubated in 30% sucrose in 0.1 M phosphate buffer overnight at 4°C until tissues sank in solution. Samples were then placed in optimal cutting temperature compound (Tissue-Tek OCT, Sakura Finetek) and embedded and frozen on dry ice. All steps were performed at room temperature unless noted.

### Blood analysis

Blood was harvested by cardiac puncture in anesthetized animals and allowed to coagulate at room temperature for 30 min to 1 hour. Blood was centrifuged at 3500 rpm for 10 min at room temperature. Serum was collected and stored at -80°C for subsequent analysis. Blood urea nitrogen (BUN) and serum creatinine analysis was carried out by Emory’s Division of Animal Resources (DAR) Diagnostic Services using ACE BUN/Urea Reagent and ACE Creatinine Reagent (Alfa Wassermann), respectively.

### H&E Histology

Paraffin-embedded kidneys were sectioned at 8 μm, rehydrated, stained with hematoxylin and eosin, and cover slipped with Cytoseal 60 (Epredia) mounting media. Kidney sections were imaged and montaged at 4x on a BioTek Lionheart FX microscope. Cyst analysis was performed in an unbiased approach using CystAnalyser software (Cordido et al., 2020). Cystic index was calculated by dividing the cystic area computed by CystAnalyser by the total kidney area computed by FIJI/ImageJ.

### Antibody and Lectin Staining

For all immunofluorescent staining, OCT-embedded tissues were sectioned at 8 μm. Tissues were rehydrated and blocked in antibody wash (1% heat inactivated goat serum, 0.1% Triton X-100 in Tris-Buffered Saline) for 1 hour. Tissues were incubated with primary antibodies overnight at 4°C, washed three times with cold antibody wash, and incubated with secondary antibodies and Hoechst 33342 for 1 hour. Slides were washed three times with cold antibody wash and coverslipped with ProLong Gold (ThermoFisher) mounting media. Slides were imaged on a BioTek Lionheart FX microscope. Primary antibodies used were: rabbit anti-ARL13B (1:500, ProteinTech, 17711-1-AP); rabbit anti-ARL3 (1:100, Kahn lab) (Cavenagh et al., 1994); rabbit anti-TULP3 (1:500, Eggenschweiler lab) (Norman et al., 2009); rabbit anti-Polycystin 2 (1:400, Stefan Somlo lab, “YCC2”) (Cai et al., 1999); rabbit anti-Cystin (1:200, Guay-Woodford lab) (Tao et al., 2009); mouse anti-acetylated alpha tubulin (1:2000, Sigma, T6793); rabbit anti-FGFR1OP (1:500, ProteinTech, 11343-1-AP); rabbit anti-NCC (1:500, StressMarq Biosciences, SPC-402D); mouse anti-THP (1:250, Santa Cruz, sc-271022). Lectins used were fluorescein conjugated *Lotus tetragonolobus* lectin (1:300, LTL, Vector Laboratories, FL-1321-2) and rhodamine conjugated *Dolichos biflorus* agglutinin (1:100, DBA, Vector Laboratories, RL-1032-2). Secondary antibodies used were goat anti-mouse AlexaFluor 647 and goat anti-rabbit AlexaFluor 488 (1:500, ThermoFisher). All steps were performed at room temperature unless noted. Cilia localization analysis of ARL3, TULP3, PC2, and cystin was performed by counting cilia using acetylated alpha tubulin and determining whether ARL3, TULP3, PC2, or cystin was detected. Counts were normalized to the cilia number in control animals so enrichment could be compared. An average of 92 cilia were counted per animal (227-323 cilia counted per stain per genotype), and three animals were counted per stain per genotype, with exception of PC2 and cystin, in which two *Arl13b*^*R79Q/R79Q*^ animals were counted (no change observed).

### Cilia length analysis

8 μm OCT-embedded kidney sections were stained with mouse anti-acetylated alpha tubulin for cilia and imaged on a BioTek Lionheart FX microscope. Z-stack images were captured at 1 μm intervals. Analysis of cilia was performed on Z-stack images using the CiliaQ plugins in Fiji/ImageJ (Hansen et al., 2021; Schindelin et al., 2012).

## Acknowledgements

We are grateful to the Eggenschwiler (Univ of Georgia), Guay-Woodford (Children’s National), Kahn (Emory) and Somlo (Yale) labs for sharing the indicated antibodies, Maya Encantada Meeks (Emory University, Division of Animal Resources) for performing blood urea nitrogen and creatinine analyses, members of the Caspary lab for discussion and manuscript comments and Quinn Eastman for editing.

## Funding

This work was supported by the National Institutes of Health: Institutional Research and Career Development Award (IRACDA) K12GM000680 and F32DK127848 to REVS; T32NS096050, diversity supplement to R01NS090029 and F31NS106755 to EDG; the Summer Undergraduate Program in Emory Renal Research (SUPERR), R25DK101390 to HGK; and R01NS090029, R35GM122549, R35GM148416, and R01DK128902 to TC.

**Supplemental Figure 1.**
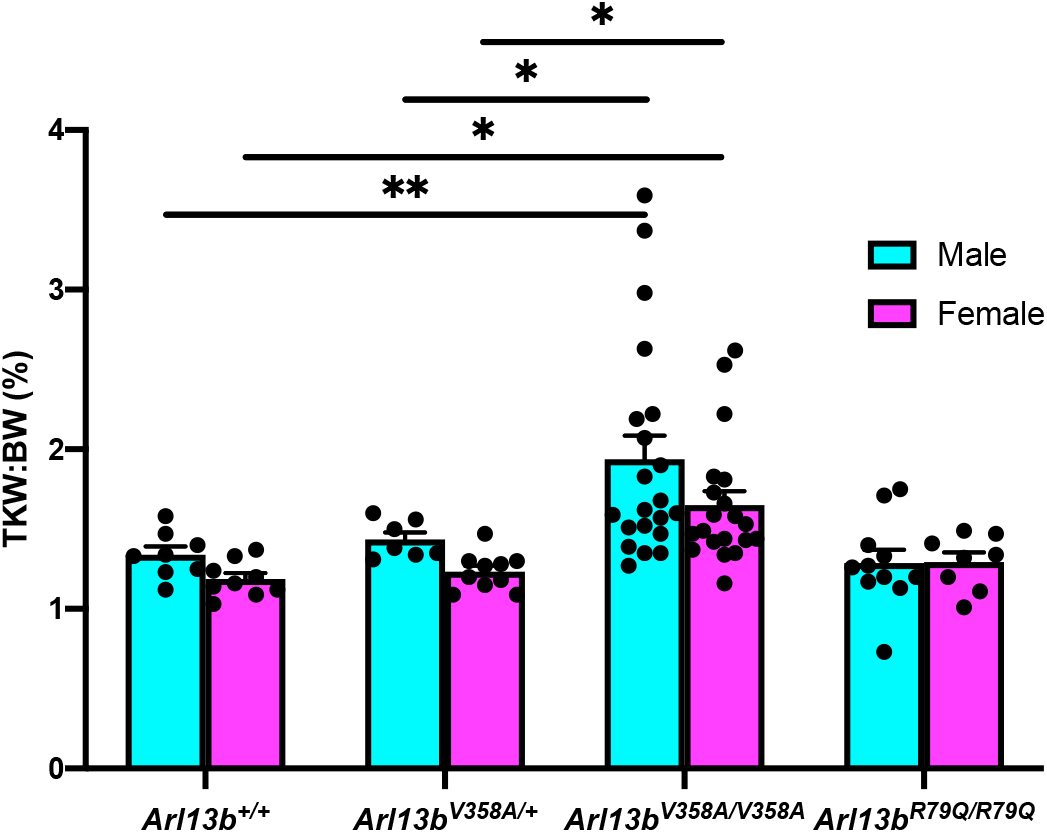
*Arl13b*^*V358A/V358A*^ mice exhibit increased kidney weight to body weight ratios. Quantification of the kidney weight to body weight ratios from 18-week-old male and female mice, mean ± SEM. Two-way ANOVA, * p < 0.05, ** p < 0.01.

