## Supplementary figures and images for "Ciliary ARL13B inhibits developmental kidney cystogenesis in mouse"

### Supplemental Figure 1

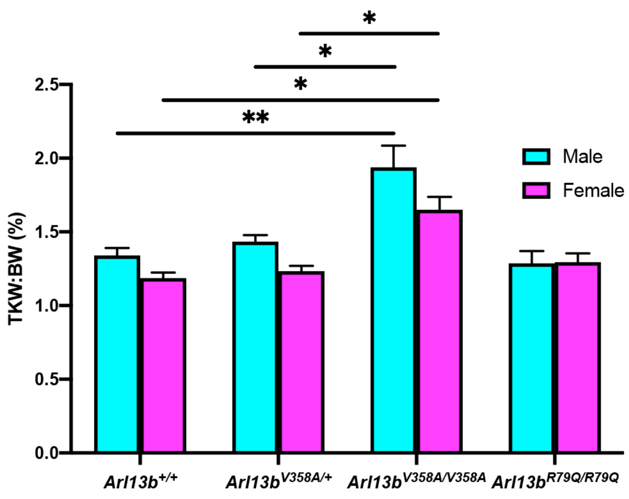
